# Multiplexing of EEG signatures for temporal and spatial distance estimates

**DOI:** 10.1101/2020.05.23.110882

**Authors:** Eva Marie Robinson, Martin Wiener

## Abstract

The perception and measurement of spatial and temporal dimensions have been widely studied. However, whether these two dimensions are processed independently is still being debated. Additionally, whether EEG components are uniquely associated with time or space, or whether they reflects a more general measure of magnitude remains unknown. While undergoing EEG, subjects traveled a randomly predetermined spatial or temporal interval and were then instructed to reproduce the interval traveled. In the task, the subject’s travel speed varied for the estimation and reproduction phases of each trial, so that one dimension could not inform the other. Behaviorally, subject performance was more variable when reproducing time than space, but overall, just as accurate; notably, behavior was not correlated between tasks. EEG data revealed during estimation the contingent negative variation (CNV) tracked the probability of the upcoming interval, regardless of dimension. However, during reproduction, the CNV exclusively oriented to the upcoming temporal interval at the start of reproduction. Further, a dissociation between relatively early frontal beta and late posterior alpha oscillations was observed for time and space reproduction, respectively. Our findings indicate that time and space are neurally separable dimensions, yet are hierarchically organized across task contexts within the CNV signal.

## Introduction

Previous research on the perception of space and time has alluded to a number of neural mechanisms responsible for the perception of each dimension (Issa, et al. 2020). However, one question that continues to be controversial is whether temporal and spatial dimensions are processed independently (Marcos & Genovesio, 2017; Cai & Connell, 2016; Robinson, et al. 2019). Previous studies have used a variety of methodologies and behavioral paradigms to provide evidence of a symmetrical or asymmetrical relationship between the perception of space and time (Bueti & Walsh, 2009; Martin, et al. 2017). Among these studies, self-motion reproduction, imagined temporal estimates, virtual navigation and temporal and spatial production have resulted in conflicting findings.

On a broader scale, areas of the brain such as the prefrontal cortex and the right parietal cortex have been implicated in different types of magnitude processing (Bueti & Walsh, 2009) especially in regard to the perception of time and space. Additional studies have provided ample evidence of the role of the right dorsolateral prefrontal cortex, supplementary motor area (SMA), basal ganglia and inferior frontal gyrus as being involved in the estimation and encoding of temporal duration through the use of fMRI (Wiener et al. 2010; Hayashi et al., 2018). Other studies have shown a neural distinction between space and time in which spatial and distance related tasks largely activate more posterior regions including the parahippocampus, anterior hippocampus, and retrosplenial complex (Gauthier & Wassenhove, 2016; Kim & Maguire, 2018; Peer et al., 2019), whereas time exclusively invokes the SMA (Coull, et al. 2015).

One well-studied neural signal associated with the measurement of temporal intervals is the slow negatively deflected Contingent Negative Variation (CNV), an event-Related Potential (ERP) component putatively driven by the SMA and corticothalamic circuitry (Nagai, et al. 2004). Previous studies on the CNV have shown a more negatively deflected pattern during different aspects of temporal processing as well as a variety of timing tasks involving subjective duration estimates. (Macar & Vidal, 2003; Monofort, Pouthas, & Ragot, 2000; Pfeuty, Ragot, & Pouthas, 2003; Ruchkin, McCalley, & Glaser, 1977). However, other experiments have called to question the specificity of the CNV for time, instead suggesting the CNV may reflect a more general neural system for different types of magnitude processing (Kononowicz & Penney, 2016; Schlichting, de Jong, & Rijn, 2017). To date, few studies have sought to investigate an asymmetry between the measurement of subjective space and time simultaneously for the CNV. One reason for the difficulty in teasing apart temporal and spatial dimensions is the natural correlation of each; longer distances naturally take more time, following time’s arrow (Riemer, 2015). Indeed, previous research has demonstrated that humans have great difficulty attending to spatial information without also considering temporal information (Glasauer at al., 2007; Kolesari & Carlson, 2017; Zach & Brugger, 2008). However, more recent studies utilizing virtual reality (VR) environments have demonstrated that time and space can effectively be decoupled from one another (Deuker, et al. 2016; Thurley & Schild, 2018; Bansal, et al. 2019; Robinson et al., 2019). In the present study, we sought to disentangle time and space processing during the estimation and reproduction of intervals from each dimension. We employed a task in which subjects walked for randomized distances in an environment lacking landmarks, crucially relying on path integration mechanisms (Wiener et al., 2016). Concurrently recorded EEG was used to measure the CNV across encoding and reproduction of both dimensions. Additionally, we spectrally decomposed EEG signals across a-priori sensors of interest (Liang, et al. 2017) in order to disentangle whether separate frequency bands contributed to distinct processing stages for each dimension. Previous studies have implicated alpha, beta, and theta frequencies during timing and spatial tasks as a form of sensorimotor integration in which theta power appears higher during movement initiation and planning in navigation tasks, indicating a possible symmetrical neural process for duration and distance perception (Bush et al., 2017, Cruikshank et al., 2012).

## Methods

### Subjects

Sixteen subjects (11 females), ages 18-35 years old from George Mason University participated in the study. Informed consent was obtained from all participants prior to the experiment and all protocols were approved by the University Institutional Review Board. All subjects were right-handed, healthy individuals without any history of neurologic or psychiatric illness.

### Task

Subjects performed temporal and spatial reproduction tasks within a virtual reality environment. The environment for the task was modified from an earlier version employed in our lab (Robinson, et al. 2019; Wiener, Michaelis, & Thompson, 2016), based on a paradigm devised by Petzschner and Glasauer (2011) and designed using the Python-based software Vizard 5.0 (Worldviz). The VR environment resembled a desert with a textured ground, 20 scattered rocks in the distance, and a clear, sunny sky. The sky was a simulated 3D dome included in Vizard software, a black and white noise image was used to create the ground texture, and a single rock was modified and imported from SketchUp 3D (Trimble Navigation) and replicated within the VR script. The construction of the VR world was such that environmental distance cues were either absent or unreliable: the initial location of the viewpoint and the position and orientation of each of the rocks was randomized at the start of every trial, and the 3D sky was such that the horizon always appeared to be a constant distance away. Participants controlled the movement of the viewpoint with a hand-held gaming controller (Xbox, Microsoft); the eye height of the VR viewpoint was set to the approximate eye height of the participant.

On a given trial, subjects were primed to walk forward, by pressing forward on the controller joystick, towards a red sphere presented on the horizon, constituting the estimation phase of the trial (Figure 1). Once moving, subjects were stopped after a particular distance had been reached. On spatial reproduction trials, this distance was selected from seven linearly-spaced intervals [4 – 10m]. On temporal reproduction trials, the distance was determined such that the time needed to reach it varied across seven linearly-spaced intervals [1 – 5s]. In either case, the speed at which the subject traveled was randomly selected from a uniform distribution between 1 and 3.6 m/s. These speeds were chosen such that the possible range of durations experienced on spatial reproduction trials matched those presented in temporal reproduction trials. After the specified distance was reached, movement of the viewpoint was stopped automatically, the horizon sphere disappeared and the environmental lighting was dimmed. The words “REPRODUCE DISTANCE/TIME” were displayed in the center of the screen. After a delay of 3s, the words disappeared, and the normal lighting resumed. Participants were allowed to simulate walking forward again (reproduction phase) and pushed a button on the controller when they judged they had reproduced the same distance or duration as during the estimation phase. Crucially, the simulated walking speed was randomly altered between the estimation and reproduction phases so that the participant could not use the time spent simulating walking as a measure of the distance traveled in distance trials, or the distance walked as a measure of time traveled in time trials. For each trial, the reproduction phase simulated walking speed was modified such that it was noticeably faster or slower than the production phase speed (maximum ±60% estimation speed, drawn from a normal distribution).

**Figure 1.**
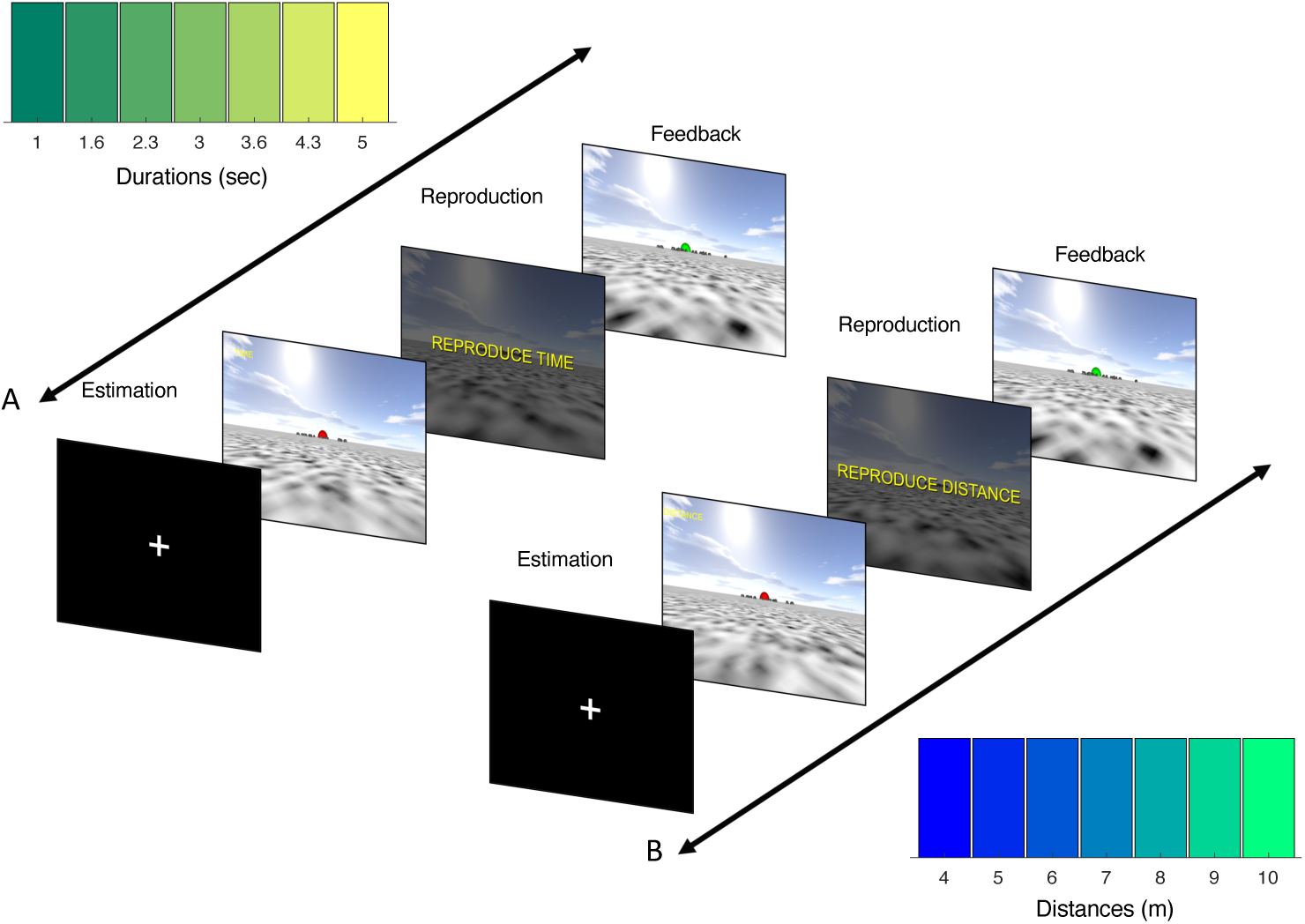
Task design schematic. Subjects performed a reproduction task with either (**A**) temporal or (**B**) spatial intervals. In both tasks, subjects were placed in an open field VR environment with no spatial cues and then initiated forward movement (Estimation). After a specified distance or time, drawn from uniform distributions (displayed), subjects were prompted to reproduce the interval they had just experienced, by again moving forward and pressing a button when they believe that interval had been reached (Reproduction). Feedback was provided on the basis of whether or not the reproduced interval fell within an adaptive window.

Following a response, subjects were provided feedback on the accuracy of their reproduction in the form of the horizon sphere, which re-appeared either as green (accurate) or red (inaccurate). Accuracy was adaptively determined on each trial by adjusting a feedback constant (*k*), such that the reproduced interval had to lie within a window [interval/*k*]; the constant was updated in a 1-up/1-down rule with a step size of 0.015. Separate values of *k* were measured for time and spatial reproduction tasks, and the initial value for each was 3.5 (Jazayeri & Shadlen, 2010).

Subjects performed the time and spatial reproduction tasks in separate blocks, with 105 trials in each block (15 trials/interval) and a single block of each. Task order was randomly counterbalanced between subjects. To analyze behavioral data, the average reproduced interval was measured for each task. In addition, we also calculated the coefficient of variation (CV), as the standard deviation of reproduced intervals divided by the tested interval. We additionally calculated the average value of the feedback constant *k* for each subject. To further analyze reproductions, we calculated the slope and intercept of a simple linear fit to reproduced intervals against the tested intervals. The slope can provide an estimate of uncertainty, as a slope closer to 1 represents veridical reproductions, whereas a slope of 0 indicates the reproduction of the same interval across all tested intervals. The intercept further provides an estimate of the offset of reproductions that can be accounted for by a general shift towards under or over-reproduction.

### EEG

EEG was recorded using an actiCHamp amplifier (Brain Products GmbH, Germany) and a 64-channel actiCAP slim active electrode montage (international 10-20 system), with FCz as the online reference and an online bandpass filter of [0.1 – 100Hz]. BrainVision Recorder (v. 1.20.0801) was used to digitize the EEG at a sampling frequency of 1000 Hz. Electrode impedances were kept below 20 kΩ. Offline processing was performed using EEGLAB for Matlab. Data were re-referenced to the mastoid channels (TP9, TP10) and down sampled to 500Hz. A low-pass Hamming windowed since finite impulse response (FIR) filter was applied at 50Hz. Electrodes with excessive noise were detected using automated methods in the *pop_rejchan* function, removed, and interpolated. The continuous EEG data was then decomposed using Infomax Independent Components Analysis (ICA); eye-blinks and other artifact components were visually determined and removed from the data. Continuous data were then epoched into separate sets for the estimation and reproduction phases of the temporal and spatial reproduction tasks, respectively. Estimation and reproduction phases were both time-locked to the initiation of movement, as this was under volitional control of the subject, in segments of [-1 – 5s].

For the examination of event-related potentials (ERPs) we focused on the CNV signal, as described above. For this, the average of a frontocentral cluster was used, consisting of electrodes [Fz, FC1, Cz, FC2, F1, C1, C2, F2, & FCz] (Wiener & Thompson, 2015). To measure significant differences between ERPs, we used the cluster-based permutation statistic by Maris & Oostenveld (2007), as implemented in Fieldtrip. We additionally chose to analyze time/frequency effects via event-related spectral perturbation (ERSP), by applying a Morlet wavelet convolution, via the *newtimef* function, by convolving a mother wavelet at 100 log-spaced frequencies between 5 and 50Hz, with 3.5 cycle wavelets and a scaling factor of 0.5. For this analysis, two clusters of interest were chosen: a frontocentral cluster corresponding to the same electrodes used for the ERP analysis, and a posterior cluster consisting of electrodes [CP1, Pz, CP2, P1, PO3, POz, PO4, P2, & CPz]; the frontocentral cluster was chosen based on our interest in beta oscillations that may underpin the CNV (Wiener et al., 2018), whereas the posterior cluster was driven by potential alpha/theta effects that may underlie spatial processing and navigation (Liang et al., 2017). Significance was again assessed via cluster-based permutation testing. Finally, we additionally examined inter-trial coherence (ITC) of phase, aligned to the onset of estimation and reproduction phases.

## Results

### Behavioral

All subjects performed the task without any difficulty. On both tasks, subjects exhibited the classic pattern of central tendency, such that the shortest interval was overestimated, and the longest interval was underestimated (Petzschner et al., 2015) (Figure 2A). Notably, this pattern was shifted slightly on the temporal reproduction task, with a greater propensity to over-estimate durations. Consistent with this finding, we also observed a significantly lower average value of the feedback constant *k* for the temporal reproduction task [*t*(15) = 3.635, *p* = 0.002, *d* = 0.91], indicating that the adaptive feedback window was wider for temporal than spatial estimates. However, no significant difference was detected between the slope of reproduced intervals for temporal and spatial reproduction tasks [Paired t-test: *t*(15) = – 0.651, *p* = 0.525], suggesting that despite the difference in veridicality, the degree of central tendency was the same; further, no correlation between slope values was observed [Pearson *r* = – 0.239, *p* = 0.374], suggesting that a common mechanism did not underlie central tendency effects observed across both tasks (Martin et al., 2017). Additionally, when analyzing CV values, a repeated measures ANOVA with task and interval as factors detected a main effect of task [*F*(1,15)=14.425, *p* = 0.002, η^2^_p_ = 0.49] but not interval [*F*(1,15)=1.079, *p* = 0.381] or interaction [*F*(1,15)=0.552, *p* = 0.768]; when collapsing across duration, this effect was shown to be due to a higher CV for the temporal reproduction task [*t*(15) = −3.798, *p* = 0.002, Cohen’s *d* = 0.95], indicating that subjects were less precise in their reproductions of temporal than spatial intervals.

**Figure 2.**
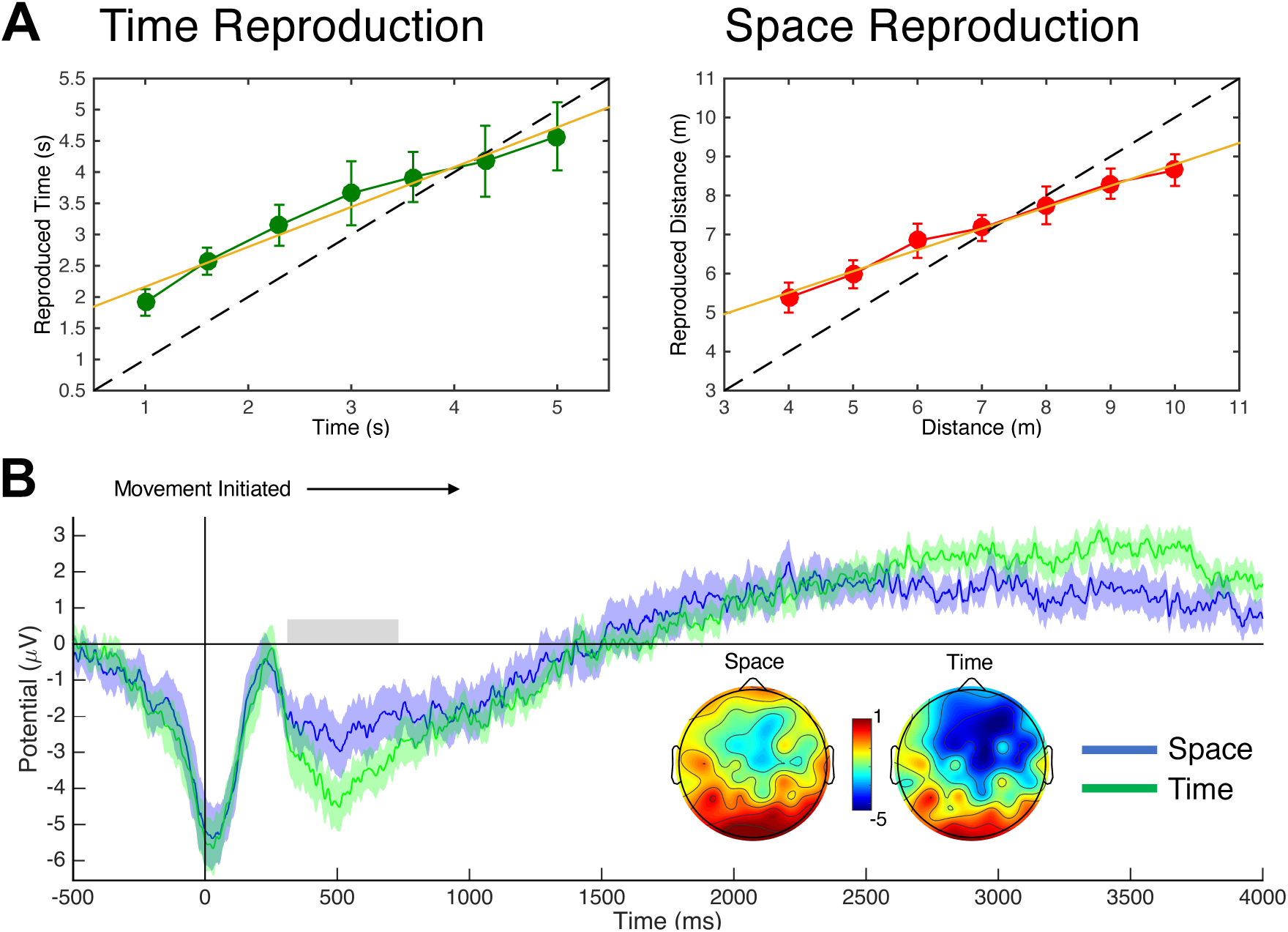
**A)** Behavioral results of the time and space reproduction tasks. Both tasks exhibited central tendency effects, with a more “centered” reproduction for spatial estimates. Linear fits (orange) exhibited no difference between tasks, whereas overall subjects were more precise for spatial than temporal estimates. Error bars represent standard error; dashed lines represent the identity. **B)** Grand-average ERPs from frontocentral electrodes time-locked to the initiation of movement in the reproduction phase. A negative deflection was observed ~500ms following movement, putatively reflecting the CNV, that was significantly larger on the temporal reproduction task; the gray bar reflects significance as determined by a non-parametric cluster test at p<0.05. Inset presents topographic maps within the significant window, demonstrating a larger frontocentral negativity for time. Shaded regions represent standard error.

### EEG

Our analysis of EEG signatures of temporal and spatial processing began by examining the CNV signal at frontocentral electrodes. For ERPs time-locked to the onset of movement initiation, we initially observed a strong negative deflection 500ms prior, reflecting the readiness potential (Brunia et al., 2012). Following this, the EEG signal rapidly returned to baseline before again exhibiting a negative peak ~500ms after movement initiation that was maximal at frontocentral sites (Figure 2B). This deflection follows the so-called “initial” CNV, or iCNV signal, which has previously been associated with attentional and arousal processes in task engagement (Tecce, 1972; Nagai et al., 2004; Fan et al., 2007), as well as time perception in particular (Kononowicz et al., 2018). Notably, the iCNV observed here was significantly larger for temporal than spatial reproductions, as determined by a non-parametric cluster test [430-598ms] (p<0.05). No other significant clusters were detected. Further, no differences were observed during the estimation phase, where both waveforms exhibited similar time courses and amplitudes (Figure 5A).

The observance of a difference in the iCNV signal between temporal and spatial reproduction tasks suggests a difference in processing between these tasks; further, the fact that this difference was observed immediately after movement initiation suggests a difference in the orienting of attention to temporal features over spatial ones (Miniussi et al., 1999; Nobre, 2001). However, one alternative possibility may relate to lower accuracy in the temporal reproduction task, which may have driven greater effort (Casini et al., 1999). To explore this difference further, we segregated the ERPs by the interval presented during the estimation phase (duration or distance). Here, we observed a striking dissociation between the iCNV signal between tasks: While the iCNV for spatial estimates remained the same, the amplitude for temporal estimates covaried with the length of the interval that the subject was about to reproduce (Figure 3). To quantify this effect, we averaged the amplitude within the window identified previously by the cluster test for each subject, and then calculated the slope of a linear regression between mean amplitude and the estimated interval. We observed that the slope of iCNV amplitudes for temporal estimates [0.68 ± 0.18 s.e.m.] was significantly different than zero [one-sample *t*-test, *t*(15) = 3.851, *p* = 0.002], whereas for spatial estimates [-0.04 ± 0.18] it was not [*t*(15)= −0.256, *p* = 0.801]; additionally, the slopes for temporal estimate amplitudes were significantly greater than for spatial estimate amplitudes [*t*(15) = 2.625, *p* = 0.019]. Interestingly, we note that the amplitudes for longer temporal intervals were associated with smaller iCNV amplitudes, a point we return to in the discussion.

**Figure 3.**
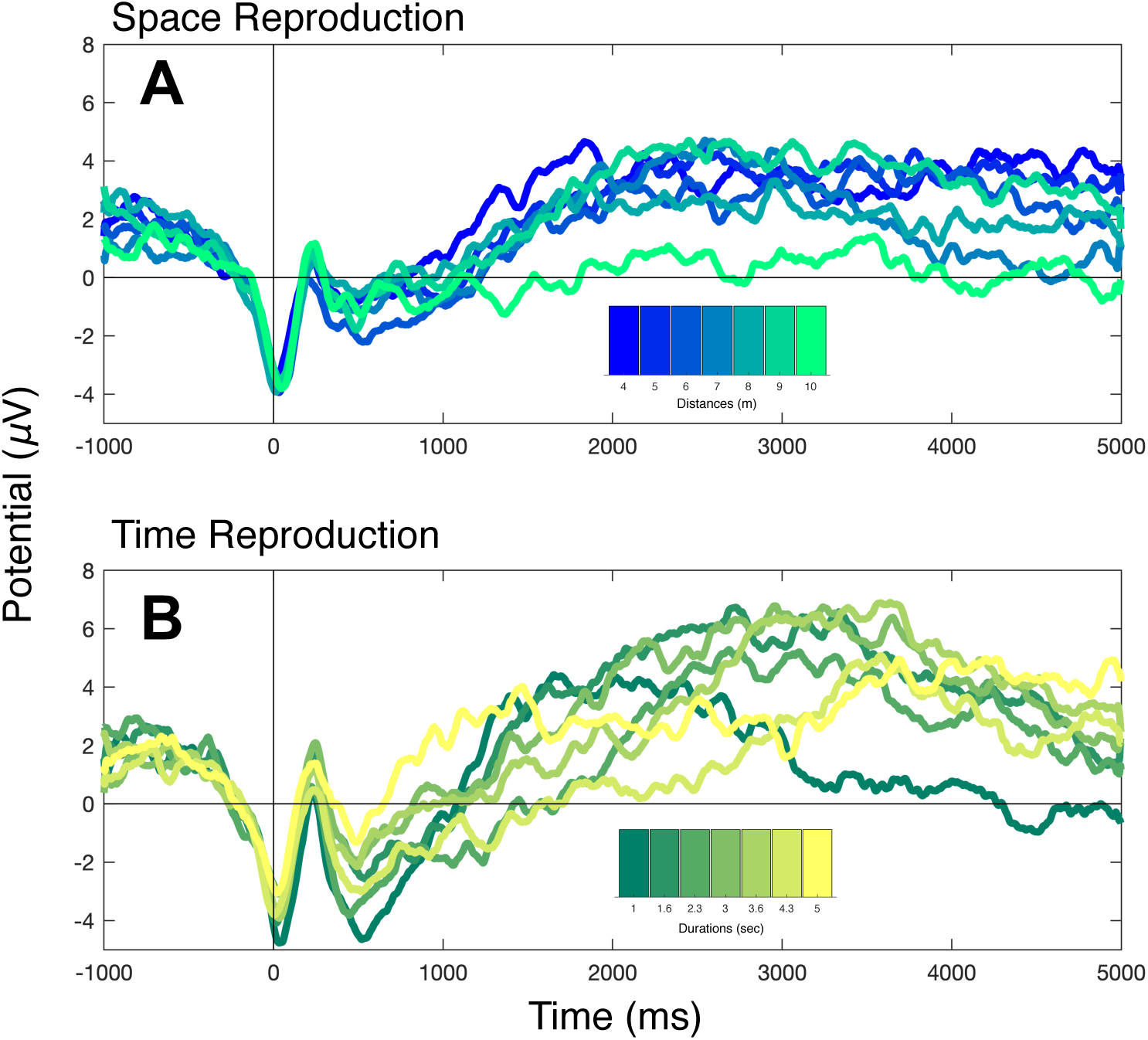
Temporal estimates change the amplitude of initial CNV. Grand average ERPs from figure 2 are presented here based on the distance (**A**) or duration (**B**) about to be reproduced. While the CNV does not differ strongly for spatial estimates, there is a linear gradient response for the CNV amplitude in temporal reproduction, with longer estimates associated with a lower amplitude [430-598ms].

For the analysis of spectral effects, we explored differences within our two a-priori regions of interest for anterior (centered on FCz) and posterior (centered on POz) electrodes. In the reproduction phase, we observed in the anterior region an increased ERSP power in a broad cluster spanning from ~7Hz to ~35Hz between ~1100ms and ~1400ms that was significantly larger for the temporal than spatial reproduction task. Within this cluster, the largest difference occurred ~15Hz, within the lower range of beta oscillations. In contrast, in the posterior cluster, an increase in power was observed for the spatial reproduction task over the temporal one in a later cluster spanning a lower frequency range (~5-14Hz) and later time (~2600-3000ms); within this cluster, the largest difference occurred ~11Hz, within the range of alpha oscillations. No other clusters were significant. Additionally, no significant differences were observed in the estimation phase.

**Figure 4.**
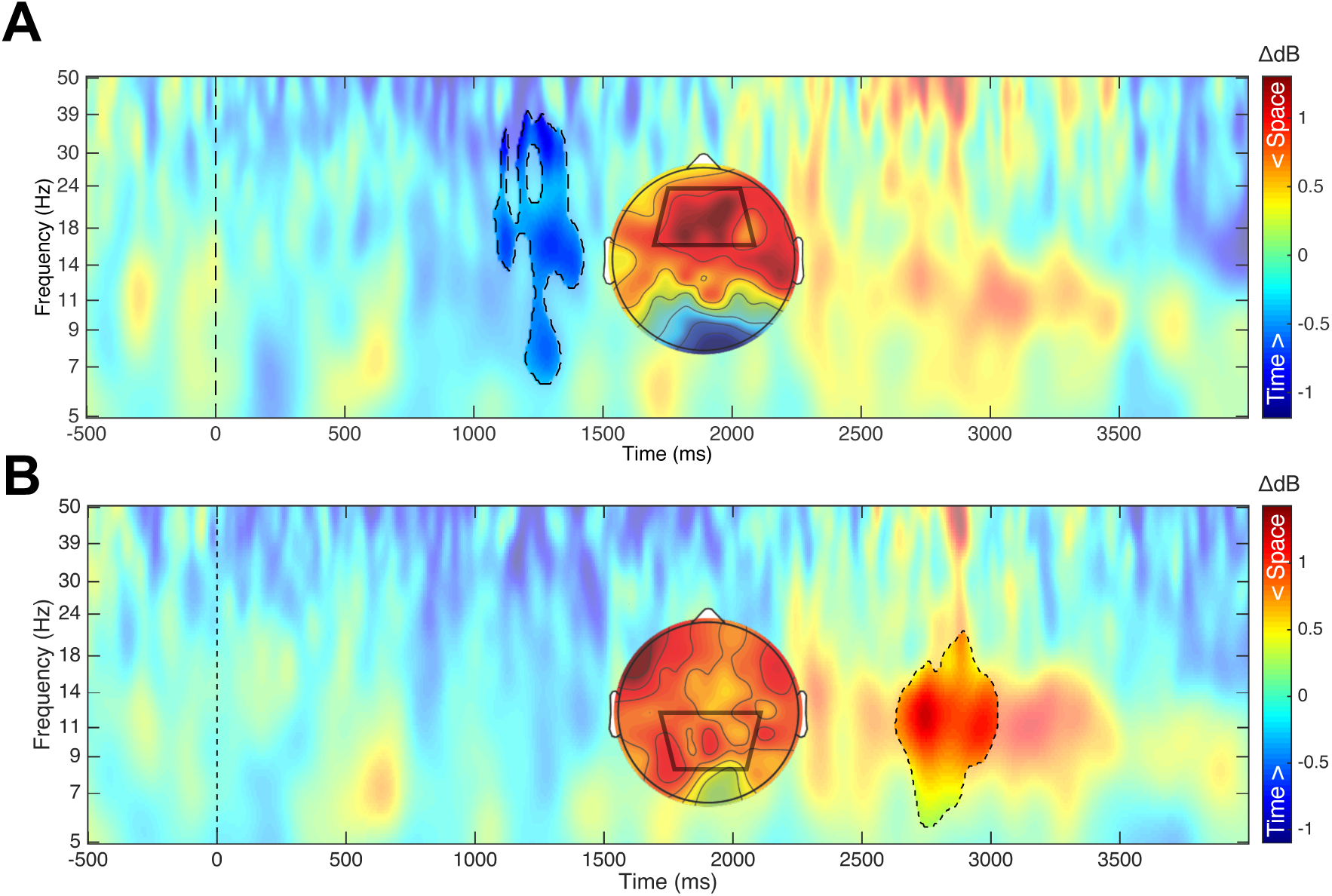
Spectral differences evoked for the reproduction phase. Both plots display ERSP results, with each pixel representing the difference between [Time – Space]. **A)** Average ERSP across frontocentral electrodes, revealing a significant cluster characterized by higher power for temporal reproduction at ~1300ms, spanning alpha to beta ranges, with a largest deflection in the low beta range (15Hz). **B)** Average ERSP across posterior electrodes, revealing a significant cluster with higher power for spatial reproduction at ~2700ms, spanning theta to alpha ranges, with a largest deflection in the alpha range (11Hz). Dashed lines indicate significant clusters. Insets represent topographic plots within the average time and frequency range of each cluster displaying power for Time **(A)** [-0.8 – 0.36 dB] and Space **(B)** [-0.54 – 0.8 dB].

For the analysis of phase synchrony across trials, via ITC, we observed an increase in synchrony in a cluster spanning ~100 – 250ms largely within the theta range (5 – 9Hz); specifically, this cluster was associated with greater coherence for temporal than spatial reproduction, and was only observed within the frontocentral cluster during the onset of the estimation phase (Figure 5). No differences were observed in either the posterior cluster or during the reproduction phase. The difference was additionally characterized by a steady increase in coherence before returning to baseline.

**Figure 5.**
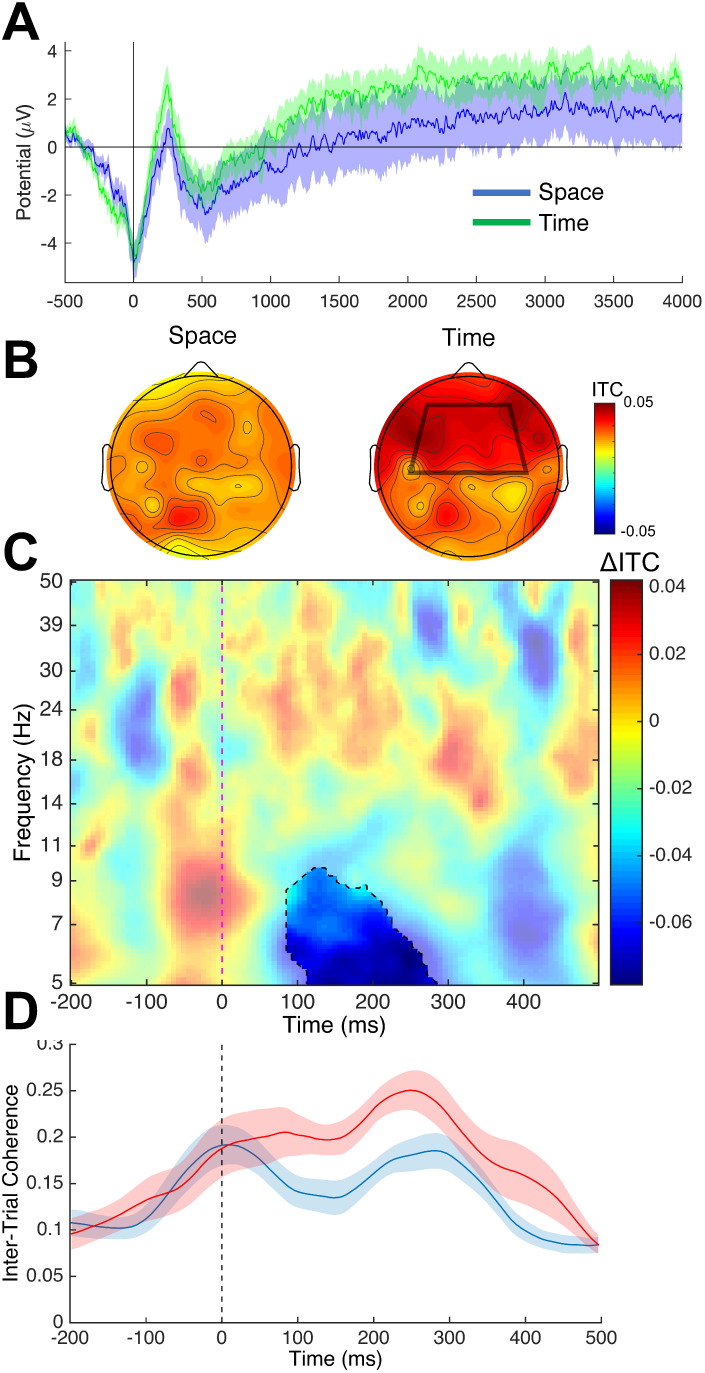
Estimation related effects. **A)** ERPs at the onset of movement initiation during the estimation phase. Both temporal and reproduction tasks exhibited similar waveforms, with no difference in the CNV or any other timepoint. **B)** Scalp topography of ITC values for FM theta from the cluster presented below. **C)** The difference in ITC values [Space – Time] time-locked to the onset of the estimation phase, averaged across frontocentral electrodes. A significant increase in ITC was observed beginning ~100ms after movement initiation spanning 5 – 9 Hz for the estimation of temporal over spatial intervals. **D)** Time course of FM theta, averaged across 5-7Hz, showing the increase in coherence corresponding to the significant cluster. Red = Time, Blue = Space. Shaded regions represent standard error.

The results of the above ERP and time/frequency analysis reveal distinct differences during the reproduction and estimation of temporal intervals. In particular, the pronounced CNV difference between temporal and spatial reproduction tasks, further distinguished by the interval-specific effects, suggest differential processing modes for the reproduction of temporal intervals. However, we note that the ERP comparison may have obfuscated any findings related to spatial, as opposed to temporal intervals. In particular, we note that in the spatial reproduction task the time needed to reach any of the distance intervals was scrambled such that any distance could be reached within a range of possible times matching the distribution used in the temporal reproduction task. As such, the time-course from onset to interval terminus is meaningless for spatial intervals, and although the CNV effects for reproduction all occurred at onset and were not distinct in terms of latency, it is possible that this difference in tasks obscured other effects.

Given the above issues, we generated a new set of ERPs, instead time-locking to the point where subjects completed the interval – either the offset during the estimation phase, or the response in the reproduction phase – so as to align each time-course to the moment when subjects completed either interval. All timeseries were mean centered, as the baseline portion was of interest (Wiener & Thompson, 2015). For the reproduction phase, we observed in both tasks a gradual negative buildup prior to the response. This signal corresponds to the so-called “late” or terminal CNV (tCNV), which underlies response preparation and anticipatory responses (Brunia et al., 2012). Notably, no differences were observed between tasks in this signal, as assessed by a cluster-based permutation test. Similarly, in the estimation phase we also observed a tCNV buildup prior to the offset, with no significant differences between tasks. However, when further separating these epochs by duration, we found a noticeable difference between estimation and reproduction phases: While in reproduction no differences were observed as a function of interval in either task, in the estimation phase we observed that the CNV prior to offset varied by the interval presented for both tasks. Here, the CNV exhibited a negative peak shift, such that the peak occurred earlier in time for longer intervals in both space and time that was also lower in amplitude (Figure 6A). This scaling of CNV responses was also observed at the single-trial level, such that the CNV stretched to accommodate either the distance or time interval to be reproduced (Figure 6B).

**Figure 6.**
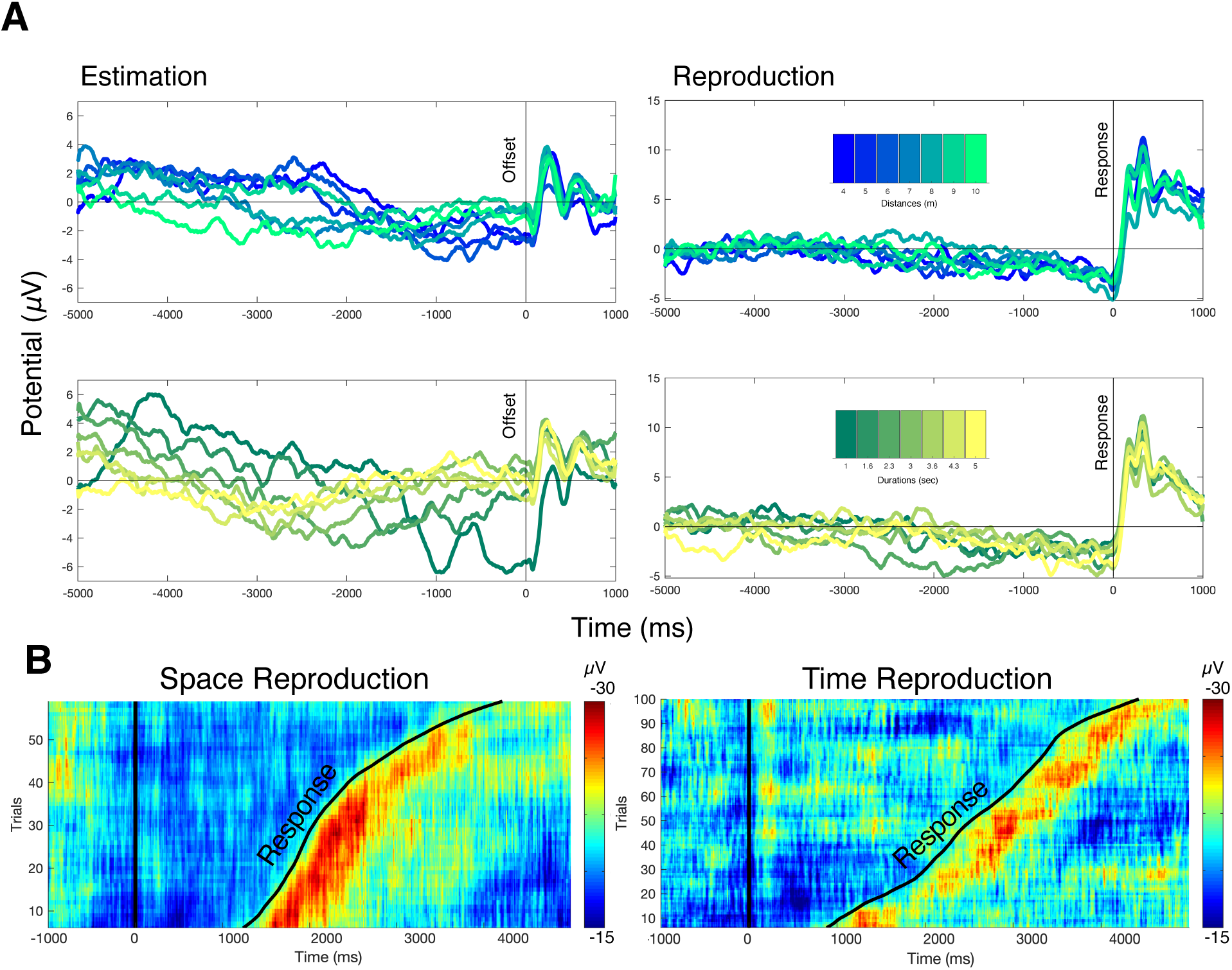
Response-locked effects. **A)** Frontocentral ERPs time-locked to either the interval offset (Estimation) or terminal response (Reproduction), divided into separate timeseries for each interval. For the reproduction phase, no difference was observed between temporal or spatial reproduction tasks or interval, with both tasks exhibiting a slow negative buildup prior to making a response to end the interval. For the estimation phase, both tasks exhibited a divergence of ERP time course, depending on the length of the estimated interval, with longer intervals exhibiting an earlier negative peak that was lower in amplitude, relative to interval offset. Critically, during the estimation phase subjects were not aware of the length of the interval being estimated ahead of time. **B)** Single trial ERP images of onset-locked reproduction data from a single subject at electrode FCz, sorted by response. In both cases, the CNV signal scaled in amplitude with the distance or time interval to be reproduced. Note that for the time reproduction image, the amplitude scaling of the iCNV at ~500ms can be observed, with more negative amplitudes for shorter reproduced intervals.

To quantify these effects, we conducted an ERP peak latency analysis, using routines provided by Liesefield (2018). Briefly, individual subject average ERPs for each interval and task were scanned for the largest negative peak deflection (100ms peak width) in the pre-onset/response window, from which the latency and amplitude were calculated. For the estimation phase, a repeated measures ANOVA on peak latencies with task and interval as within-subject factors found a linear main effect of interval [*F*(1,14)=8.277, *p* = 0.012, η^2^_p_ = 0.372] and a task by interval interaction [*F*(1,14)=16.39, *p* = 0.001, η^2^_p_ = 0.539], but no effect of task [*F*(1,14)=0.444, *p* = 0.516], characterized by a greater linear dispersion of peak latencies in the duration task. Similarly, for peak amplitude, a linear effect of interval [*F*(1,14)=11.844, *p* = 0.004, η^2^_p_ = 0.458] and task by interval interaction [*F*(1,14)=5.509, *p* = 0.034, η^2^_p_ = 0.282], but no effect of task [*F*(1,14)=1.606, *p* = 0.226] was observed in which peak amplitudes decreased with increasing interval magnitude and a larger difference for time than for space reproduction. For the reproduction phase, no significant effects were observed for peak latency or amplitude across any interval or task (all *p* > 0.05).

## Discussion

In the current study, subjects completed a temporal and spatial reproduction task within a virtual reality environment, presented on a desktop computer screen. In this study, we analyzed EEG signatures first by examining the CNV at frontocentral electrodes and found a strong negative deflection prior to the onset of movement, followed by a strong negative deflection after movement initiation. The initial negative deflection, or iCNV, which has been associated with time perception, attention and arousal during task engagement, was significantly larger for temporal reproduction than spatial reproduction. As previously noted, the temporal and spatial reproduction tasks exhibited a difference in the iCNV signal, suggesting that subjects needed to orient their attention to temporal features over spatial features. (Liu et a. 2013; Taatgen, van Rijn & Anderson, 2007).

Critically, there was a dissociation between the iCNV signal between tasks where the amplitude for temporal estimates covaried with the upcoming reproduced interval, while no difference in the iCNV for spatial estimates was observed. We also note that the amplitudes for longer temporal intervals were associated with a smaller iCNV signal. This finding has been reported in previous studies. Specifically, Kononowicz and Colleagues (2015) used a temporal discrimination task to investigate the hypothesis that iCNV and late CNV amplitudes reflect the accumulation of temporal pulses over time and found that longer temporal reproductions were associated with smaller CNV amplitudes. A central framework of timing continues to be the pacemaker-accumulator model in which the brain contains a pacemaker whose pulses are integrated by an accumulator to process measures of time.

Included within this theory is the notion of climbing of neural activity over time that is associated with the SMA and is indexed by the CNV. Interestingly, our results and the findings by Kononowicz and colleagues are in contrast to the assumptions of climbing neural activity, where the accumulation of temporal information should be directly related to a larger increase in CNV amplitude. Instead, our results point to a scaling of the CNV signal for both space and time. While these findings may seem counterintuitive, they accord with recent electrophysiological recordings from rodents (Matell, et al. 2011) and monkeys (Renoult, et al. 2006; Lebedev, et al. 2008; Jazayeri & Shadlen, 2015) demonstrating that firing rates scale to match the temporal interval. Further, a recently developed firing-rate theory of time perception posits that an accumulation regime can be characterized as opponent firing rate processes between neuronal populations that also scale to the temporal estimate (Simen, et al. 2011). Consistent with this theory, recent work has shown that, at the onset of a reproduction interval, the amplitude of population firing (Jazayeri & Shadlen, 2015), as well as the overall dynamic configuration of that network (Wang, et al. 2018; Remington, et al. 2018), vary with the length of the interval. Indeed, our own findings of iCNV amplitude scaling support this “pre-planning” hypothesis, in which the to-be-reproduced interval is set in advance.

Yet, while the reproduction CNV signal demonstrated clear dissociations between tasks, the estimation CNV did not. Indeed, here we observed that the CNV signal again scaled with the measured interval, such that the peak in amplitude was linearly shifted earlier in time from the interval offset, regardless of time or distance. Importantly, subjects were not aware of the interval they were measuring on any given trial. This finding accords with recent work on anticipation, in which the probability of the interval offset grows with the elapsed interval, mathematically expressed as a hazard ratio (Nobre, et al. 2007). Previous work has demonstrated that the CNV signal is sensitive to the probability of upcoming events and can be an index of the hazard ratio (Scheibe, et al. 2009; Mento & Vallesi, 2016; Breska & Deoull, 2017; Boettcher, et al. 2020). However, all of these studies have shifted upcoming events in time; in our present study, we found that the CNV can also account for the elapsed distance as well. Crucially, as time was scrambled across possible distances by randomly varying the speed gain, the CNV estimates of distance probability cannot be accounted for simply by elapsed time.

These finding suggests a multiplexing of the CNV signal across phases and task requirements; during measurement, anticipatory responses can be oriented separately in space and time, whereas during reproduction, temporal estimates can be set in advance. Yet, why are distance estimates not also pre-planned? While it remains possible that a pre-planning signal for distance exists, but was not observed in the present study, an alternative possibility is that pre-planning was not crucially necessary for distance reproduction. Indeed, path integration mechanisms, in the absence of any viable landmarks, may critically rely on accumulating information retrospectively from optic flow regions (Chrastil & Huang, 2019; Zajac, et al. 2019). However, when landmarks, even distal ones, are present, the brain may utilize these for prospectively orienting to the goal location (Brown, et al. 2016; Ekstrom, et al. 2017). We further suggest that, during estimation, subjects prospectively orient themselves in both space and time to future possible intervals, but that during reproduction, subjects are oriented prospectively in time, and retrospectively in space.

the iCNV amplitude may not be an ideal representation of temporal measurement and may instead largely reflect a change in processing, specifically an orienting and anticipatory response (Bender et al. 2004; Fischer et al. 2008, Fischer et al. 2010). Additionally, the iCNV signal observed was significantly higher for temporal reproductions than spatial reproductions, suggesting that while the iCNV may underlie an orienting response, it may be reflective of an index of measurement that is unique to timing. This assertation is also supported by the observed dissociation between the iCNV between the temporal reproduction task and the spatial reproduction task.

Furthermore, our results suggest an asymmetric and possible disentangled relationship between the perception of spatial and temporal information in that measuring intervals of time and space require different attentional processes and therefore, rely on separate neural systems (Brunec et al., 2017; Jafarpour & Spiers, 2016; Robinson et al., 2019). Our analysis of spectral effects further these conclusions. Here, we observed a significantly larger ERSP power increase in a broad cluster spanning from late theta to late beta (7Hz – 35Hz) for temporal reproductions over spatial reproductions. This difference was seen within the anterior region of interest and was largest for low beta. In contrast, an increase is alpha power specifically was seen for the spatial reproduction task in the posterior region of interest. More generally, these findings provide continuing support for the involvement of theta, alpha and beta oscillations in the measurement of temporal and spatial intervals (Bischof and Boulanger, 2003; Hsieh, Ekstrom, & Ranganath, 2011; Liang, Starrett, & Ekstrom, 2018; Samaha et al., 2015; Vilhelmsen et al., 2015; Kononowicz & Rijn, 2015; Kulashekhar, Pekkola, Palva, & Palva, 2016; Javadi, et al. 2019). More specifically, they also offer a window into distinct processing stages for each task type; increased frontal beta was observed relatively early in reproduction (~1300ms), shortly after the first possible interval (1000ms) could have elapsed, whereas increased posterior alpha was observed relatively late in reproduction (~2700ms), towards the probable end of most distances. One possibility here is that frontal beta oscillations underlie proximity to a memory standard used for judging relative duration (Wiener, et al. 2018; Mendoza, et al. 2018), whereas posterior alpha oscillations underlie the encoding of spatial distances in memory (Khader, et al. 2010; Sauseng, et al. 2005; Sutterer, et al. 2019). Notably, in an fMRI version of the distance reproduction task used here, greater hippocampal activation was observed as subjects completed distance reproductions, which further correlated with the degree of central tendency between subjects (Wiener, et al. 2016).

While both beta and theta frequencies being active for timing and spatial tasks has been thought to reflect a symmetrical neural process for these measurements, our findings, combined with ERP analysis reveal distinct differences in neural underpinnings for the processing of temporal and spatial information with different phases of the CNV being associated with different cognitive aspects of measuring time or space. It is important to note that the current study as well as additional research on the relationship between the CNV and time perception has provided evidence that the CNV may be better understood as representing several subcomponents that encompass different cognitive aspects of measuring time.

Overall, our findings support a multiplexing of EEG signatures during temporal and spatial processing. Under this framework, the CNV generally indexes the probability of spatial and temporal intervals, but specifically the planning of reproduced times. Further, oscillatory phase and amplitude separately index the relative temporal and spatial position, while reproducing those intervals, across theta, alpha, and beta bands. These results broadly support separate yet overlapping processing regimes for time and space, with distinct networks and operating modes for each stimulus dimension.

## Notes

Conflict of Interest: The authors declare no competing financial interests.

### Competing Interest Statement

The authors have declared no competing interest.

